# Whole-brain MEG decoding of symbolic and non-symbolic number stimuli reveals primarily format-dependent representations

**DOI:** 10.1101/731687

**Authors:** Brett B. Bankson, Daniel Janini, Chris I. Baker

## Abstract

The human brain can rapidly form representations of numerical magnitude, whether presented with symbolic stimuli like digits and words or non-symbolic stimuli like dot displays. Little is known about the relative time course of these symbolic and non-symbolic number representations. We investigated the emergence of number representations for three stimulus formats - digits, words, and dot arrays - by applying multivariate pattern analysis to MEG recordings from 22 participants. We first conducted within-format classification to identify the time course by which individual numbers can be decoded from the MEG signal. Peak classification accuracy for individual numbers in all three formats occurred around 110 ms after stimulus onset. Next, we used between-format classification to determine the time course of shared number representations between stimulus formats. Classification accuracy between formats was much weaker than within format classification, but it was also significant at early time points, around 100 ms for both digit / dot and digit / word comparisons. We then used representational similarity analysis to determine if we could explain variance in the MEG representational geometry using two models: a GIST feature model capturing low-level visual properties and an approximate number model capturing the numerical magnitude of the stimuli. Model RSA results differed between stimulus formats: while the GIST model explained unique variance from 100-300 ms for all number formats, the performance of the approximate number model differed between formats. Together, these results are consistent with the view that distinct, format-specific number representations, moreso than a single “abstract” number representation, form the basis of numerical comparison.

## Introduction

The human brain can support a multitude of different representations for number. These representations enable both the estimation of the number of objects in our environment and formal mathematics over number symbols like digits. When the brain receives sensory input from a set of objects, it represents their numerosity through an approximate number system (ANS) (Feigenson et al., 2004). This representational system is shared among many animals including prelinguistic human infants (Xu and Spelke, 2000), monkeys (Cantlon and Brannon, 2006), crows (Ditz and Nieder, 2015), and fish (Agrillo et al., 2012, Piffer et al., 2013). In addition to this phylogenetically ancient system of representation, modern literate humans also represent number through written symbols of digits and number words. The extent to which these symbolic and nonsymbolic number representations rely on shared neural substrates has been queried for decades. These efforts have primarily focused on whether the same brain areas implement symbolic and nonsymbolic number representations, while fewer studies have compared the time course of symbolic and nonsymbolic number representations. In order to address the ways in which symbolic and nonsymbolic number representations rely on shared versus distinct neural resources, we must address both when and where these representations are implemented. In the current study, we coupled magnetoencephalography (MEG) with multivariate decoding and representational similarity analysis (RSA) to elucidate the temporal dynamics of number processing across distinct representational formats.

Extensive neuroscientific evidence supports the view that approximate number representations are implemented by neural populations within parietal and frontal cortex. A key hallmark of the ANS is its relationship to Weber’s law such that the discriminability of two sets of objects depends on their ratio rather than their respective absolute values (Feigenson et al, 2004). Neuroscientific work in both non-human primates and humans has revealed analogous neural tuning for number in lateral prefrontal cortex and intraparietal sulcus (Piazza et al., 2004, Bulthé et al., 2014, Nieder, 2016), supporting the view that these regions form the basis of the ANS.

In order for visual symbols like digits and number words to activate numerical representations, they must first be categorized. This process is putatively achieved by the reading circuits of the ventral visual pathway (Dehaene, 2009), culminating in the formation of a number form or word form representation tolerant to low-level changes in the font, size, and position of the visual symbol. Within this system, there is ongoing debate surrounding the extent to which the formation of number and word forms depends on shared or distinct neural regions within the ventral visual stream (Yeo et al., 2017). After a visual number symbol is categorized, representations of its meaning can be activated. A central question in the study of numerical cognition is what these symbolic number representations entail. One possibility is that number symbols activate the same representations as nonsymbolic dot displays, more specifically the ANS. An alternative possibility is that number symbols primarily gain numerical content by activating representations distinct from the ANS, perhaps concepts involved in abstract logic and language rather than concepts that ground out in visual perception.

Although it is unclear how these symbolic and nonsymbolic number representations are implemented neurally, behavioral experiments indicate that nonsymbolic and symbolic number are partially represented though shared resources. For instance, participant reaction times when comparing the magnitudes of two digits are a function of numerical distance between the digits, suggesting the use of an analog scale similar to the ANS (Moyer and Landauer, 1967; Dehaene et al., 1990). There has been extensive debate about whether number symbols activate neural populations in parietal cortex that represent “abstract” number, meaning number representations elicited by symbolic and non-symbolic number stimuli across sensory modalities (Kadosh and Walsh, 2009). Lussier and Cantlon (2017) recorded fMRI activity as participants compared the magnitudes of numbers and found that the level of activity in the intraparietal sulcus is modulated by numerical ratio for both symbolic and nonsymbolic number stimuli. Moreover, it has been reported that intraparietal sulcus adapts to repeated presentations of the same magnitude for both digits and dot displays, and that recovery from this adaptation can occur across these stimulus formats (Piazza et al., 2007). In contrast, multivoxel pattern analyses suggest that symbolic and nonsymbolic number representations are implemented by different patterns of activation within intraparietal sulcus (Bulthé et al., 2014; Bulthé et al., 2015). Thus, while behavior indicates a link between nonsymbolic and symbolic number representations, it is less clear how this link is instantiated by the brain despite the large number of studies investigating the spatial localization of number representations.

While most prior work has focused on the spatial localization of number representation, there is a relative lack of understanding of the time course of these representations. Recently, Teichmann et al. (2018) used MEG and multivariate pattern analyses to study how neural representations of symbolic number (digits and dice) emerge over time. Their findings suggested that format-specific representations of symbolic number emerge within 150 ms of stimulus presentation, and more tentatively that shared representations between the two symbolic formats emerged later around 400 ms after stimulus presentation. Here, we build upon these findings by investigating the time course of both symbolic and nonsymbolic number representations rather than just symbolic number representations. Using MEG, we measured the neural response to visual number stimuli (values 6-13) in the following formats: 1) digits, 2) number words, 3) and dot displays. We used a decoding approach to determine how quickly the brain forms representations of individual numbers within each of these formats. Next, we determined whether we could find evidence of shared number representations by conducting cross-decoding across formats. Finally, we used RSA to determine when models of low-level visual shape and number magnitude predicted the neural responses in the brain. By emphasizing the temporal dynamics of visual number processing, we offer new means of comparing the neural substrates that underlie symbolic and non-symbolic processing.

## Methods

### Participants

22 healthy participants (17 female, age range 20-44) with normal or corrected-to-normal vision participated in the current study. All participants gave written informed consent before participation as a part of the study protocol 93-M-0170, NCT00001360. This study was conducted according to the Declaration of Helsinki and was approved by the Institutional Review Board of the National Institutes of Health.

### Stimuli

We created three sets of number stimuli that ranged from 4-18 in magnitude (Figure 1). One set contained numbers represented as digits, a second set contained numbers represented as words, and third set contained numbers represented as dot arrays. These three sets allowed us to examine visual processing of symbolic (digits, words) and non-symbolic (dots) number formats. All three stimulus sets were presented in white, subtending a maximum of 6° x 6° of visual angle and centered on a black background (participant viewing distance: 70 cm). To maximize within-format variability in visual features, 32 unique exemplars were generated for each magnitude in the digit and word stimulus sets. 26 of these exemplars were formed from different fonts, and the other 6 exemplars were formed using hand-written scripts from 3 individuals who were not involved with the study. A similar procedure was used for the dot array stimuli, whereby 32 unique exemplars for each number were generated with a script by Gebuis and Reynvoet (2011).

**Figure 1.**
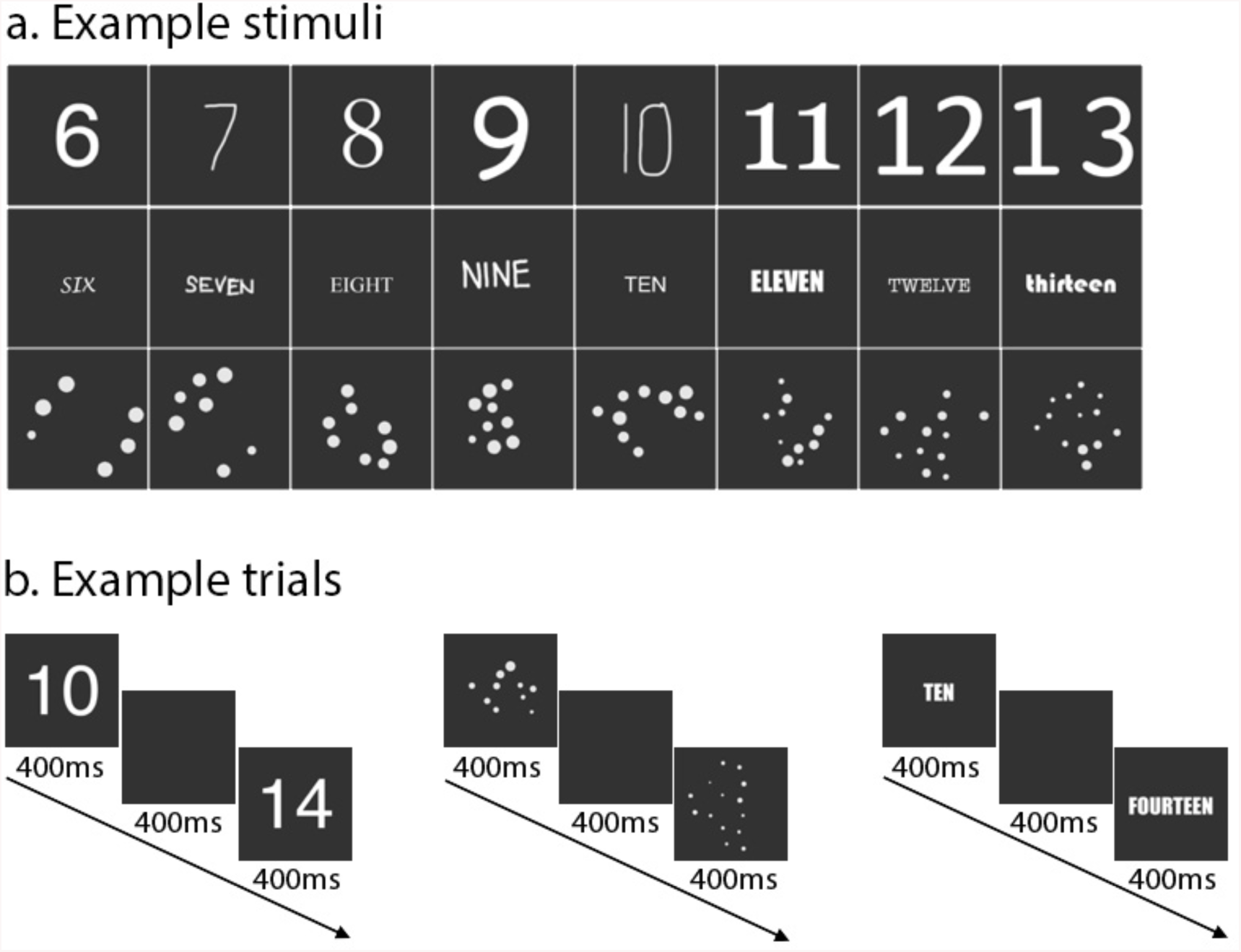
Example stimuli and trial progression. **a.** 32 different stimuli were generated for each number in each format. Here we show one example for each number in each format. **b.** Stimuli were presented on a black background for 400ms, followed by a blank black screen for 400ms, and then followed by the second stimulus for 400ms. Upon presentation of the second stimulus, participants judged whether the second stimulus was larger or smaller than the first stimulus and responded via button press. The first stimulus was always a number from 6-13.

### Procedure

For the MEG recordings, participants entered an electromagnetically shielded MEG chamber where they were seated upright within the dewar. Stimuli were presented with the Psychophysics Toolbox (Brainard, 1997) in MATLAB (version 2016a, Mathworks, Natick, MA). Visual presentation was controlled by a Panasonic PT-D3500U DLP projector with an ET-DLE400 lens, located outside the chamber and projected through a series of mirrors onto a back-projection screen in front of the seated participant.

### Task

Participants completed a magnitude comparison task during MEG recording. While fixating, participants were presented with a number for 400 ms, followed by a delay period with blank screen of 400 ms, a second number for 400 ms, followed by an inter-trial interval of 1800 ± 100 ms that consisted of a blank screen and fixation cross. Participants responded after the presentation of the second number with a button press to indicate whether the second number was larger or smaller than the first number. The first number was always between 6 and 13, and the second number was always 20% or 40% smaller or larger than the magnitude of the first number, rounding to the nearest whole number. Because discriminability of number magnitudes is a function of the number pair ratio, we controlled for task difficulty by maintaining a set ratio between number pairs in this task.

One complication with number comparison over dot displays is that many visual cues also tend to increase along with numerosity. The script used to generate our dot-stimuli (Gebuis and Reynvoet, 2011) accounted for this potential confound by minimizing the extent to which the visual cues of area extended, density, surface area, item size, and circumference predict numerical distance between pairs of numbers. Thus, participants had to encode the actual numerosity of the dot display stimuli in order to complete the task rather than simply attending to one of these other visual cues.

Participants completed 12 experimental runs that were divided into 4 blocks of 3 runs each, with self-paced breaks between each block. Each run contained 128 trials with a fixed number stimulus format, i.e. only digits, or words, or dot arrays. Within each format, participants were presented with each number 6-13 a total of 64 times. Each run lasted 384 seconds, resulting in a total experimental time of 76 minutes.

### MEG acquisition and preprocessing

MEG data were recorded continuously with a 275-channel CTF whole-head MEG system at a sampling rate of 1200 Hz (MEG International Services, Ltd., Coquitlam, BC, Canada). All analyses were conducted in MATLAB (version 2017a, The Mathworks, Natick, MA). Preprocessing steps used Brainstorm 3.4 (version 02/2016, Tadel et al., 2011) and custom-written code similar to recently published MEG decoding work (Bankson et al., 2018). Recordings were obtained from 272 channels (dead channels: MLF25, MRF43, MRO13), consisting of radial first-order gradiometer channels with synthetic third-gradient balancing to remove background noise online. Participants’ head position was localized at the beginning of the experiment and after each experimental block, using fiducial coil readings at the nasion, left and right preauricular points. We recorded this head position information to provide feedback about the quality of head placement in the dewar. Data were bandpass filtered between 0.1 and 300 Hz, and bandstop filtered at 60 Hz and harmonics. Data were segmented into single trial bins consisting of 100 ms pre-stimulus baseline activity for normalization purposes and 900 ms activity after the first number presentation of each trial.

To increase SNR and decrease computational load, we employed three additional pre-processing steps (outlined in Bankson et al, 2018): PCA dimensionality reduction, temporal smoothing on PCA components, and data downsampling. Principal components analysis (PCA) was run to reduce the number of channels into the set of most descriptive components. All data for an MEG channel across trials were concatenated for PCA, and the components explaining the least variance were removed to speed-up further processing, with a maximum removal of 30% of the components (i.e. 80 components) or 1 % of the variance, whichever was reached first (Hebart et al., 2018). For all participants, the smallest 80 components explained less than 1% of the variance, so the data for all further analyses contained 192 components. Data across all time points were normalized according to the baseline period of −100 to 0 ms relative to stimulus presentation. To do so, the mean and standard deviation of the baseline period for each component were computed, and the mean was subtracted from the data before dividing by the standard deviation. We then used a Gaussian kernel of ± 15 ms half duration at half maximum (HDHM) to temporally smooth the remaining components, and downsampled the components to 120 Hz (121 samples / trial).

### Multivariate decoding and cross-classification

We first used time-resolved multivariate classification of MEG data within each participant to examine the representational dynamics of symbolic and non-symbolic number stimuli. To determine the extent to which distributed neural representations for different numbers are discriminable from one another over time, we used a linear support vector machine implemented with LIBSVM in MATLAB (SVM; Chang & Lin, 2011). Analyses are based on general guidelines for multivariate MEG analysis (Grootswagers et al., 2017). Functions from The Decoding Toolbox (Hebart et al., 2015) and custom written code were used for subsequent analyses, which were applied to all participants.

Because our stimuli comprised both symbolic (digits and words) and non-symbolic (dots) number stimuli, we focused our analyses on identifying the emergence of discriminable representations of individual numbers both within and across stimulus formats. Below, we outline analyses for within-format pairwise classification and between-format pairwise cross-classification. This set of analyses allowed us to investigate the possibility of format-specific and format-independent representations of number.

### Within-format SVM classification

The following within-format classification steps were conducted independently for each stimulus format of digit, word, and dot array trials. For each format, we created supertrials in training and test sets by averaging 4 trials of the same number drawn randomly without replacement (Isik et al., 2013). At every downsampled time point, preprocessed MEG data for each supertrial were arranged as a *P* dimensional vector (equal to the number of components from PCA preprocessing). This yielded *K* pattern vectors for each time point and number. For each pair of numbers at every time point, we used leave-one-out classification by training a classifier on *K*-1 pattern vectors and testing on the pair of left-out pattern vectors.

The random generation of supertrials and subsequent classification procedure of assigning training and testing sets was repeated 100 times for each pair of numerosities at each time point. The resulting decoding accuracies were averaged across the 100 iterations and yielded an 8 x 8 matrix at every time point, with the rows and columns indexed according to numbers 6-13 and the diagonal left undefined. To evaluate average pairwise decoding accuracy, we computed the average of the lower triangular matrix (excluding the diagonal).

We assessed significance for the within-format decoding analysis with a sign permutation test. We ran the decoding procedure 1,000 times for each participant, then randomly multiplied the resulting accuracy values within each iteration by +1 or −1. These sign-permuted accuracies were averaged across all participants to generate a null distribution of decoding accuracies. *P*-values were determined as one minus the percentile rank of the veridical group mean in this null distribution. These *p*-values were corrected according to the false-discovery rate (FDR) and were considered significant if the corrected *p*-value did not exceed 0.05 in a one-tailed test and was contiguous with at least 2 other significant time points.

### Between-format SVM cross-classification

The following between-format classification steps were conducted between digit and word trials, digit and dot array trials, and word and dot array trials. Cross-classification used the same preprocessing steps as within-format classification. At each time point for each pair of numerosities, we trained a classifier on all supertrials in format 1 and tested this model on all supertrials in format 2. This was repeated by training on format 2 and testing on format 1, and the whole process repeated 100 times with different supertrial assignment each time. Because training and testing data were extracted from independent experimental runs, all supertrials within a given classification permutation were included as opposed to using leave-one-out classification. Pairwise accuracy values in the form of an 8 x 8 matrix for both directions of classification were averaged together to yield an average cross-format classification result. Average cross-format pairwise decoding accuracy was evaluated by computing the average of the lower triangular matrix, with the diagonal defined in this case. Significance was assessed for the between-format cross-classification procedure using the same sign permutation test steps as outlined above for the within-format classification.

### Representational similarity analysis

RSA allows the comparison of neural signals and predictive models by abstracting patterns of information from modality-specific representations (Kriegeskorte et al., 2008). In this study, we were interested in comparing the neural representational space with two models: a GIST feature model capturing low-level visual properties of each stimulus and a second model based on the number magnitude represented by the stimuli. We converted MEG patterns into representational dissimilarity matrices (RDMs) that quantify the pairwise relationship between all patterns of experimental conditions. At each time point, we quantified how much variance in the MEG RDM was accounted for by each model.

### MEG similarity matrices

To construct RDMs from the MEG data, we first computed the pattern of response elicited by each number at each time point. We calculated the mean pattern of response in the preprocessed 192-component space for all trials of each number. This yielded 8 MEG patterns (one for each number 6-13) for each of the three formats at each time point. Within each format, we used a Spearman correlation to compute the similarity between all pairs of the 8 patterns, and subtracted these correlation values from 1 to result in three 8 x 8 MEG RDMs for each time point. These RDMs were analyzed further by quantifying their relationship with model matrices, as described below.

### Representational dissimilarity matrices for GIST features and approximate number

To characterize the temporal evolution of number-related information in the MEG signal, we compared two models to MEG data: a GIST feature model that provides an account of gross visual differences between stimuli, and an approximate number model based on the properties of the ANS.

The GIST model describes the distributions of orientations and spatial frequencies present in the stimuli (Oliva and Torralba, 2001). Each image was passed through a bank of Gabor filters with 3 spatial frequencies and 12 orientations for high spatial frequencies, 8 orientations for moderate spatial frequencies, and 6 orientations for low spatial frequencies (26 filters). Filter outputs were computed in an 8 x 8 grid, resulting in 1664 features. We computed the pattern of response across these features for each stimulus. A Spearman correlation was computed between all pattern vectors within a format, yielding a 256 x 256 meta-matrix. This matrix was subtracted from 1 to generate a dissimilarity matrix. We computed the mean dissimilarity across the 32 exemplars per number to yield an 8 x 8 RDM for each format.

We generated the approximate number RDM from the pairwise dissimilarities in log-scaled magnitude of all numbers 6-13. By using the log-transform of absolute pairwise differences, we more closely approximate the tuning curves of the ANS that have been shown to govern number representations outside of subitizing range (numbers 1-4). This model was equivalent for all three number formats.

### RDM comparisons

We first computed the correlation of our models to assess their general similarity, before comparing them to MEG signal. Spearman’s *r* was calculated for each pair of models, and the significance of correlations was tested with a row shuffled randomization test: for the pair of models in question, the rows and columns of the first RDM were randomly permuted before computing the Spearman’s *r* between the second model RDM. We repeated this procedure 1,000 times to generate a null distribution of correlation coefficients, and the results were judged to be significant if they showed a higher correlation coefficient than the distribution cut-off determined at *p* < 0.05.

### Variance Partitioning: Unique and Shared Contributions

Given that our two models could explain overlapping portions of the variance in the MEG RDMs, we conducted a variance partitioning analysis to determine the unique and shared variance accounted for by each model (see Groen et al., 2012, Greene et al., 2016, and Bankson et al., 2018, for similar approaches). We accounted for the variance in the MEG RSMs using different combinations of RDMs as regressors: 1) a ‘complete’ regression with each RDM serving as a predictor, 2) a ‘single-predictor regression’ with only the GIST RDM as a predictor, and 3) another ‘single-predictor regression’ with only the approximate number RDM as a predictor. We subtracted the explained variance (*R*^*2*^) values of these different regression analyses to measure the partitions of variance uniquely explained by each model, and the variance explained by both RDMs. We determined statistical significance by running a row shuffled randomization test as described above: rows and columns of model matrices were randomized 1000 times and the original analysis repeated. The same randomization index was used across all models to match the randomization test assumptions, and the significance cutoffs for *R*^*2*^ values were set to *p* < 0.01 (FDR-corrected) and required to be contiguous with at least 2 other significant time points. Because these statistical analyses are permutation based, they implicitly test against the baseline of variance rather than an alternate null hypothesis of *R*^*2*^ = 0. We established a variance baseline by repeating the above variance partitioning analysis with two noise models and simulated MEG data (all generated from random number assignment) to demonstrate the non-zero variance baseline.

## Results

### Temporal dynamics of within-format number representations

To quantify the time course of representations for individual numbers, we used time-resolved multivariate decoding and conducted pairwise classification between MEG signal patterns in response to number stimuli in digit, dot array, or word formats (Figure 2). Pairwise classification was conducted only for MEG signal in response to the first number presented in each trial. Individual digits could be differentiated rapidly after stimulus onset, peaking at 110 ms (mean accuracy: 75.04%) and showed a slow decay in decoding accuracy that remained significantly above chance for the majority of the first stimulus trial window (800 ms). Individual words showed a similar time course but lower decoding accuracy, peaking at 110 ms (63.4%) and remaining significantly above chance until ∼600 ms after stimulus onset. Individual dot arrays again showed a similar peak in decoding accuracy at 110 ms (57.55%) but had less sustained decoding accuracy than the other two stimulus formats. These results indicate that neural representations of number arise quickly regardless of presentation format. However, these representations could be format-specific or could be shared across formats. To test the nature of the representations, we next conducted cross-decoding between formats.

**Figure 2.**
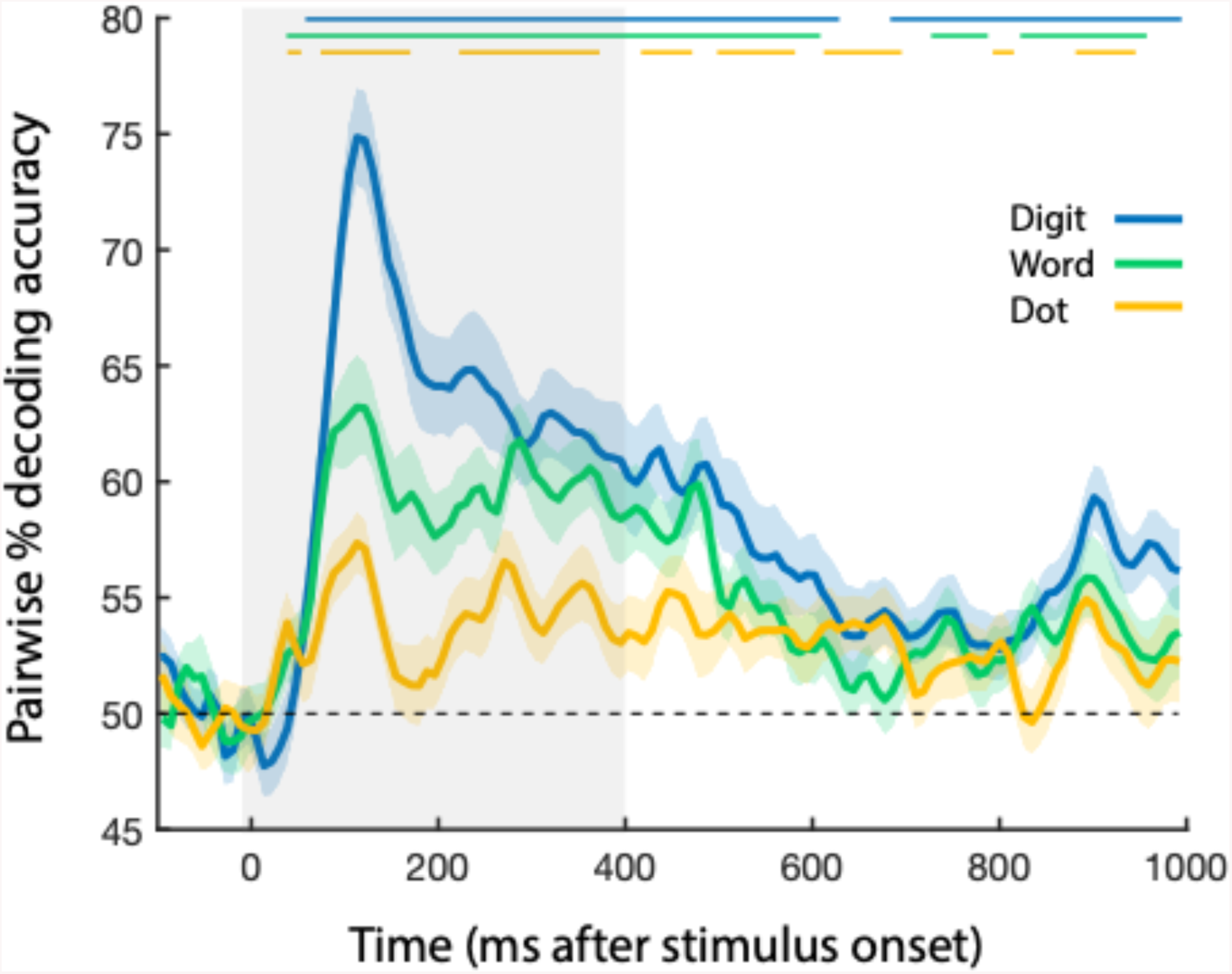
Time-resolved within-format number classification. After the onset of the object stimulus (depicted in gray from 0-400 ms), pairwise number classification accuracy increased rapidly for all three number formats. Error bars reflect SEM across participants for each time point separately. Significance is marked at the top of the figure, corresponding to *p* < .05 (FDR corrected).

### Temporal dynamics of between-format number individuation

We trained a linear SVM classifier on one format then tested it on another, completing this process for all pairs of formats: digits and words, digits and dots, and words and dots. This procedure was conducted in both directions, and the results averaged (i.e. train on dots, test on digits; train on digits, test on dots). Digits and dots showed a first peak at 100 ms (mean accuracy: 52.34%), with significant above chance classification accuracy from 60-120 ms and at several later time points between 290-685 ms after stimulus onset. Digits and words showed a similar early peak at 100 ms (52.06%) and a second peak at 290 ms (52.15%); digit and word cross-classification was significantly above chance from 40-110 ms and 270-370 ms after stimulus onset. Word and dot classification was never significantly above chance. The results here suggest shared number representations that are more limited than within-format number information. These shared number representations also exist to a greater degree between digits / dots and digits / words than dots / words in the context of this magnitude judgment task.

### Model Similarity

We compared MEG signal to two models: a GIST visual feature model and an approximate number model. To quantify the relationships between the models derived from GIST features and number magnitude, we computed the correlation between the model RDMs (Figure 4a). GIST and number magnitude models were most strongly correlated for dot array stimuli (*r* = 0.72, *p* < .001), followed by digit (*r =* 0.39, *p* = 0.02), and word stimuli (*r* = 0.26, *p* = 0.18). The high correlation between GIST features and number magnitude for dot array and the modest correlation for digit stimuli suggests that number decoding within and between these formats may be driven by GIST features as opposed to associated magnitude information. Because of these significant correlations, we conducted variance partitioning analyses to determine how much unique variance number magnitude versus GIST features could account for in the MEG signal.

**Figure 3.**
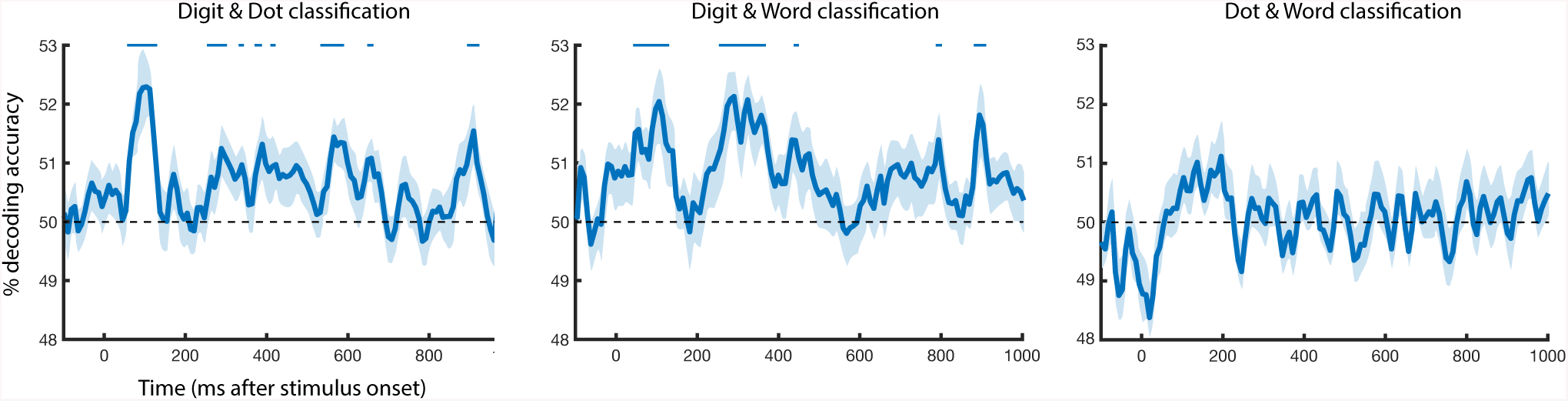
Time-resolved between-format cross classification. Classifiers were trained and tested on digit and dot stimuli, digit and word stimuli, and dot and word stimuli to ascertain representational overlap between different number formats at each time point. Pairwise cross-classification accuracy increased after stimulus onset for digit / dot and digit / word comparisons, but not for dot / word comparison. Error bars reflect SEM across participants for each time point separately. Significance is marked at the top of the figure, corresponding to *p* < .05 (FDR corrected).

**Figure 4:**
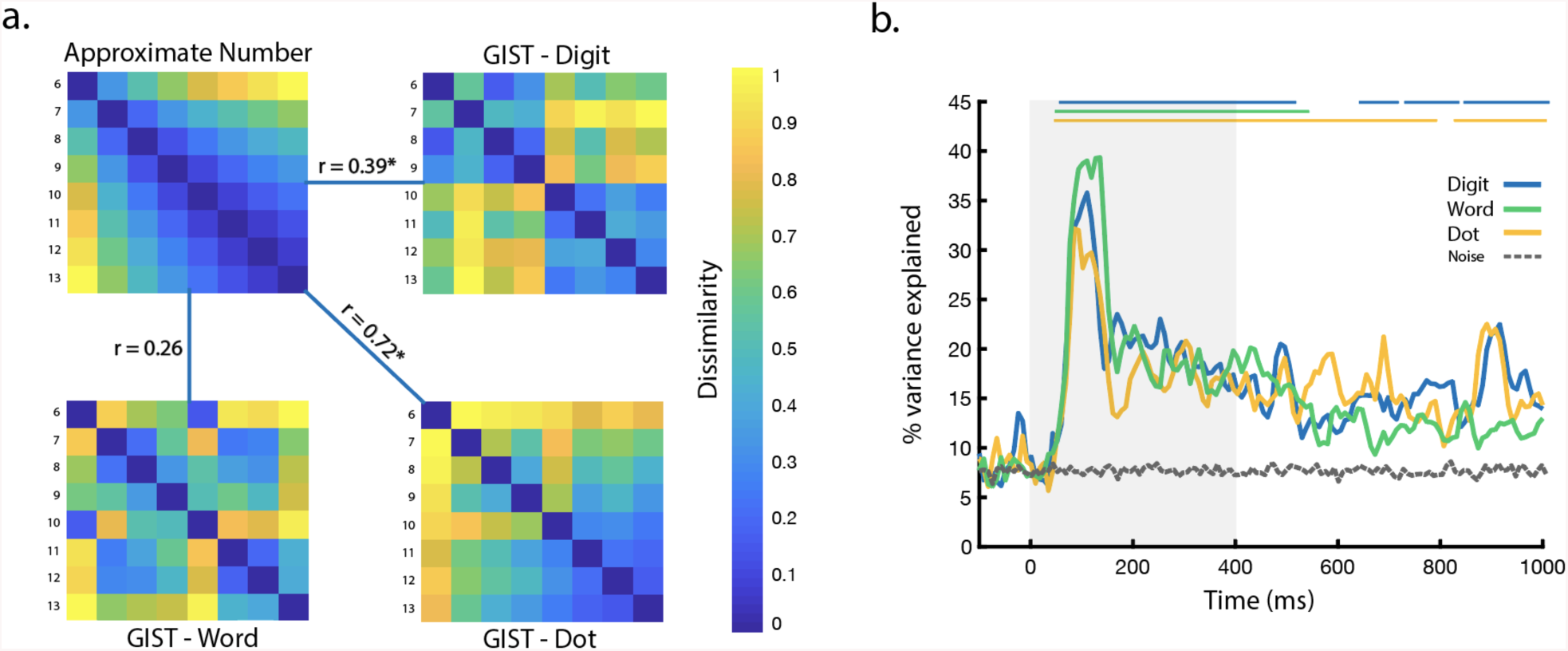
**a**. RDM comparisons between approximate number model with format-specific GIST models for digit, dot, and word stimuli. RDMs are plotted by rank to enhance visual contrast. Spearman correlations between the approximate number model and GIST for digits and dot stimuli were significant (*p* < .05), assessed with row permutation tests. **b.** Time resolved variance partitioning showing the total variance explained by both models for MEG signal in response to each stimulus format. Baseline variance is plotted in gray. Significant time points are marked at the top of the figure, corresponding to *p* < .01 (FDR corrected).

### Variance Partitioning

We conducted a variance partitioning analysis that described the unique variance in the MEG response accounted for by each model and the shared variance accounted for by both models. We used a threshold of *p* < 0.01 (FDR-corrected) to determine significant model contributions to MEG variance (Figure 4b).

For MEG responses to digit stimuli, the GIST model explained unique variance at early time points, 70-370 ms after stimulus onset with a peak at 110 ms (*R*^*2*^: 23.5%) (Figure 5). In contrast, the approximate number model explained significant portions of MEG variance primarily after 760 ms, with a peak after presentation of the second stimulus at 916 ms (*R*^*2*^: 18.9%). The GIST model and approximate number model accounted for shared variance from 85 – 270 ms after stimulus presentation with a peak at 110 ms (*R*^*2*^: 7.8%). The total variance explained by unique and shared model contributions was significant from 70 – 520 ms and 630 – 1000 ms after stimulus presentation with a peak at 110 ms (*R*^*2*^: 35.8%).

**Figure 5.**
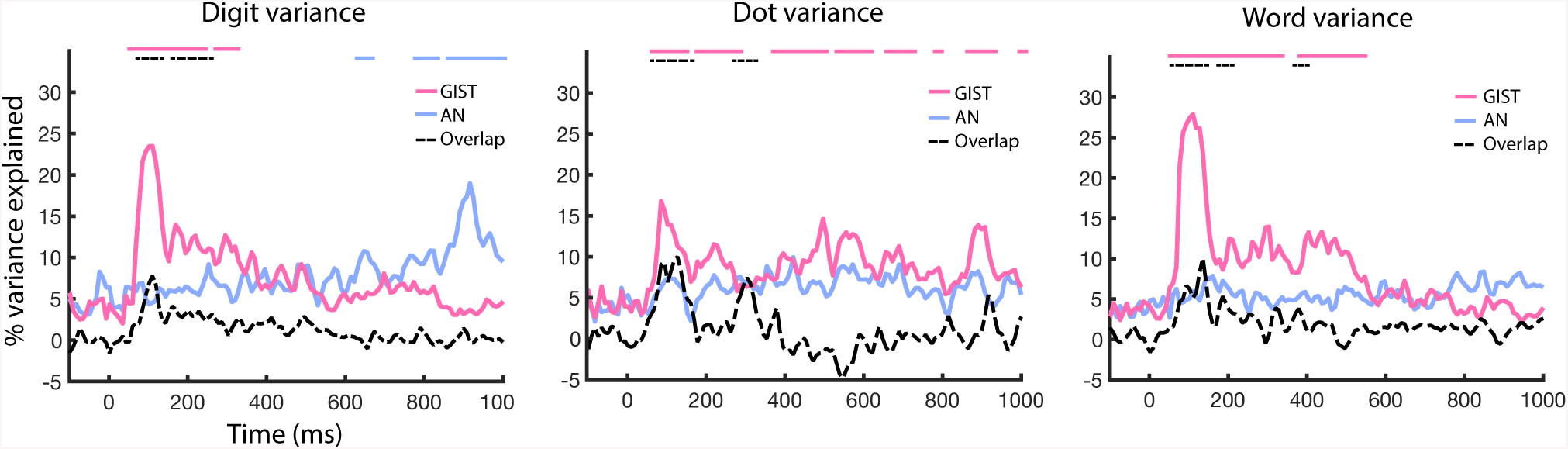
Time resolved variance partitioning showing the total unique and shared variance explained by both models for MEG signal in response to each stimulus format. Significant time points are marked at the top of the figure, corresponding to *p* < .01 (FDR corrected).

For MEG responses to the dot array stimuli, the GIST model explained significant variance from 75 – 260 ms and again sporadically between 390 – 980 ms after stimulus onset, with a peak at 85 ms (*R*^*2*^: 16.8%). The approximate number model did not significantly explain any unique MEG variance throughout the entire time course. The combination of GIST + approximate number models explained a significant portion of the variance from 75 – 160 ms (peak 150 ms, *R*^*2*^: 10.1%), and then between 280 – 320 ms. The slightly negative deflection of shared variance between GIST + approximate number models from 400-600 ms is not atypical for variance partitioning analyses: this pattern suggests that the GIST model does not capture information that is relevant to the approximate number model, and vice versa (Pedhazur, 1997). The total variance explained from unique and shared model contributions was significant from 60 – 790 ms and again from 820-1000 ms, with a peak at 85 ms (*R*^*2*^: 32.15%).

Finally, for MEG responses to the word stimuli, the GIST model explained unique variance 70-350 ms after stimulus presentation and later from 390-510 ms, with a peak at 110 ms (*R*^*2*^: 27.9%). The approximate number model did not significantly account for any unique MEG variance during the entire time course. The GIST model and approximate number model explained shared variance starting at 70 ms after stimulus onset until 230 ms (peak 130 ms, *R*^*2*^: 10.1%), and briefly from 350-390 ms. The total variance explained by unique and shared model contributions was significant from 60-520 ms after stimulus onset, with a peak at 135 ms (*R*^*2*^: 39.4%).

## Discussion

In this study, we examined the time course of number representation in both symbolic and non-symbolic formats from patterns of whole-brain MEG signal. Our results support the existence of both distinct and shared representations for symbolic and non-symbolic number. Using within-format decoding, we show that individual digits, number words, and dot arrays can all be classified above chance within 110 ms of stimulus presentation. This suggests that format-specific representations of digits, words, and dot displays have similar temporal dynamics, emerging early after image presentation, then persisting throughout the trial. Using between-format classification, we demonstrated shared representations between digits & words and digits & dots at early (60-110 ms) and later (300-450 ms) latencies. Finally, model-based RSA showed predominant contribution of the GIST model to early MEG variance in response to all number formats, whereas an approximate number model explained significant variance solely for symbolic digit MEG responses at longer latencies.

Our results on within-format number classification indicate that representations for individual numbers can be accurately decoded within the first 100 ms of stimulus presentation, regardless of symbolic or non-symbolic format. Using a similar paradigm, Teichmann et al. (2018) also reported above chance classification for individual numbers presented as digits or dice, though in their study significant classification emerged later in time. While significant classification in our study emerged at ∼50ms after stimulus onset and peaked at 110ms, significant classification in their study emerged around 120-145 ms and peaked at 250ms. Importantly, we did not vary the retinotopic position of our stimuli, while Teichmann et al. did. The early representations reported in our experiment may be retinotopically specified, whereas the later representations reported in Teichmann et al. may be tolerant to variation in retinotopic position. Support for this claim comes from the fact that the early representations in our study were well explained by the GIST model. This pattern of results is consistent with previous MEG studies indicating that the earliest time points following stimulus presentation carry retinotopically specific representations, whereas position-invariant representations begin to emerge by about 150 ms (Wardle et al., 2016; Isik et al., 2013).

Our cross-classification results between digits and word stimuli provide some evidence of shared representations between symbolic number formats. Previously, Teichmann et al. (2018) also reported evidence for shared representations between two symbolic formats: digit and dice stimuli. They show significant between-format decoding for a brief period around 400 ms, suggesting a late emergence of a shared number representation. Similarly, we found limited evidence of shared representations from 300-400 ms, but we also demonstrated significant cross-classification between digits and words at very early time points from ∼50 – 110 ms after stimulus presentation. Our results suggest that associations between symbolic formats might be an early component of the visual representation for number. In both studies, this association may be due to shared word representations between the two stimulus formats rather than shared magnitude representations.

Our cross-classification results between digits and dot displays suggest that associations between digit representations and the magnitude representations of the ANS may arise within 100 ms of stimulus onset. Our stimulus set (numbers 6-13) was chosen to avoid numbers in or near the subitizing range, so all non-symbolic numbers were represented by the ANS rather than working memory systems that rely on parallel individuation. Therefore, the association between symbolic and non-symbolic number in our study is likely supported by the ANS. These results are consistent with behavioral findings that adults can accurately compare symbolic and non-symbolic number up to about the number twelve, though the associative mapping is weaker for higher numbers (Sullivan and Barner, 2013). In contrast to our study, Teichmann et al. (2018) utilized the numbers 1-6, so most of their stimuli were nameable numbers within the subitizing range. By using larger numbers outside of this range, our findings build upon these previous results and provide some evidence that the association between digit representations and the ANS is registered automatically and quickly by the visual system.

Although the dot / word cross-classification did not yield any periods of significant decoding, this null result cannot speak to the existence or lack of representational overlap between number words and dot stimuli. These two formats showed the weakest within-format classification accuracies, so perhaps a higher-powered study focusing just on these two formats would yield the data necessary to investigate whether shared representations can be found between number words and the ANS.

The variance partitioning analyses allowed us to tease apart when the GIST model and an approximate number model explained variance in the neural representations for number stimuli. For digits, the MEG signal contains an early response within the first 100 ms that is uniquely explained by the GIST model as well as shared information between GIST + approximate number models to a certain degree. Later, the MEG signal for digits is increasingly explained by the approximate number model rather than the GIST model. Strikingly, the approximate number model explains the most variance at 916 ms, or 116 ms after the presentation of the second number stimuli in each trial. This latency precisely coincides with the timing by which magnitude information from the first stimulus becomes behaviorally relevant. This pattern of results suggests that neural responses transitioned from representing visual information to representing magnitude information at the time in the trial when those magnitude representations became task-relevant. Results from the word stimuli showed contrasting results: the approximate number model did not explain the MEG signal across the entire time course, and instead the GIST model uniquely explains a majority of the variance within 500 ms of stimulus presentation. The unique pattern of results for digits in comparison to words could indicate the frequency and facility with which we manipulate number information in the form of digits as opposed to number words.

The model analyses for the dot stimuli highlight the unavoidable fact that approximate number representations are highly correlated with other low-level visual features. The correlation between the GIST model and the approximate number model for the dot displays was r = 0.72 while it was much smaller for the digit and word displays (r = 0.39 and r = 0.26, respectively). This exemplifies that the mapping between visual features and numerosity is fairly arbitrary for number symbols, but highly meaningful for dot arrays: GIST features provide dominant information on number magnitude. Neuroscientific experiments (Harvey et al., 2013; Nieder et al., 2002) and psychophysical experiments (Burr and Ross, 2008; Cicchini et al., 2016) have established that number representations can be formed without the use of any one low-level visual feature that usually correlates with number. At the same time, accuracy of number comparison is affected by the ways in which low-level visual features covary with number, suggesting that the visual system ordinarily relies on these low-level visual features to form more accurate number representations (Gebuis et al., 2009; Gebuis and Reynvoet, 2012). Future MEG decoding studies could address the current observations by systematically varying low-level features of dot display stimuli to explore their role in tuning dynamic representations of approximate number.

Despite the fine-grained temporal resolution of our analyses, we cannot comment on the spatial origin of the representations being studied here. Particularly with regards to the early contributions of the GIST model to MEG signal variance across all three number formats, an important expansion of this work could entail using human intracranial recordings to examine the spatial extent of early visual activity in representing symbolic and non-symbolic number across ventral temporal and lateral parietal areas.

Many studies have searched for shared representations between symbolic and non-symbolic number with the assumption that these “abstract” number representations provide the foundation for mathematical cognition (Gallistel and Gelman, 2000; Dehaene, 2007; Dehaene, 2009; Piazza, 2011). We agree that ANS representations play a role in some everyday mathematical tasks; the heavily replicated distance effect supports the view that the ANS plays a role in common number comparison tasks in both children (Holloway and Ansari, 2009) and adults (Libertus et al., 2007; Moyer and Landauer, 1967; Dehaene et al., 1990). Moreover, structural alignment processes may allow the ANS to be recruited broadly when reasoning about any magnitude, for example reward probabilities (Luyckx, 2019). However, neither empirical evidence nor theoretical arguments support the view that the ANS is the primary foundation of mathematical cognition. While individual studies have argued for stronger effects, a recent meta-analysis concluded that the ability to compare the magnitude of non-symbolic number stimuli is only weakly correlated with mathematical achievement (r = 0.241, CI [.198, .284]) (Schneider et al., 2017). More importantly though, a primary source of mathematical thought during development is the construction of integer representations when learning to count, and these representations cannot in principle be supported by the ANS (Carey, 2009). Adults and children alike can form integer representations that exactly enumerate sets, giving us the knowledge that 278 is exactly one less than 279; the *approximate* number system is by its very definition incapable of supporting this knowledge. In order to understand how mathematical thought gets off the ground, we not only need to understand how number symbols are associated with ANS representations, but also how the brain forms exact representations that transcend the limitations of the ANS. Representations unique to symbolic number play a foundational role in mathematical thought, a role that could never be filled by “abstract” number representations shared for digits and dot displays.

Collectively, our results provide evidence that representations of numerosity and number symbols are formed from dot displays, digits, and number words within 100ms after stimulus presentation. These representations are largely format-specific as evidenced by 1) higher decoding accuracy for within-format classification as compared to between-format classification, and 2) heterogeneous model contributions to the MEG signal for each stimulus format. We do find some evidence for shared representations between symbolic and non-symbolic number at early (∼100 ms) and later (∼300 ms) timepoints, though evidence for a robust and singular “abstract” number representation was much weaker than evidence for format-specific representations. Our results support the view that multiple format-specific representations, more so than a singular “abstract” number representation, underlie the ability to compare numerical magnitudes. In order to more fully understand the neural underpinnings of mathematical thought, future work will need to characterize how the brain implements integer representations in a symbolic number system in concert with approximate number representations in a nonsymbolic number system.

## Acknowledgments

This work was supported by the Intramural Research Program of the National Institute of Mental Health (ZIA-MH-002909) – National Institute of Mental Health Clinical Study Protocol 93-M-0170, NCT00001360.

## Notes

Conflict of Interest: The authors declare no competing financial interests.

